# Revisiting daily growth and survival insights of Southern bluefin tuna (*Thunnus maccoyii)* larvae in the eastern Indian Ocean

**DOI:** 10.1101/2025.09.10.675289

**Authors:** R. Borrego-Santos, J.M. Quintanilla, R. Laiz-Carrión, A. García, E. Malca, F. Abascal, D. Die, I. Riveiro, R. Swalethorp, M.R. Landry

## Abstract

This study analyzed the growth patterns and survival of Southern bluefin tuna (SBT, *Thunnus maccoyii*) larvae collected during January–February 2022 in their only known spawning area in the eastern Indian Ocean (IO). Otolith microstructure was examined to characterize both population-level and intra-population growth (OPT-optimal and DEF-deficient group), with special emphasis on the flexion process, as well as to provide insights into larval survival. SBT larvae began flexion at sizes and ages comparable to those reported in other bluefin tuna species. At the intra-population level, larvae reached flexion at the same age; however, optimal (OPT) larvae reached this stage in better physical condition, with greater length, weight, and body depth, likely increasing their chances of survival at later stages. The observed larval growth rates (0.38 mm·d⁻¹) exceeded those from a historical study in 1987 (0.33 mm·d⁻¹), likely due to a ∼2 °C increase in sea surface temperature and shifts in prey availability. Larval survival appears to depend on a selective process based on growth, in which only a small proportion of individuals (<2%) exhibited width increment in otoliths similar to those of surviving larvae, allowing for faster development and earlier access to larger prey. These findings highlight the need for expanded research on the early life stages of SBT, particularly in the context of ongoing ocean warming and climate change.

## 1. Introduction

Southern bluefin tuna (*Thunnus maccoyii*, SBT) is an important scombrid species for tuna fisheries and a commercially valuable resource for countries around de Indian Ocean (IO) (Waples et al., 2018; Macfadyen, 2019). The population declined to critically endangered status in the 2000s (Anon, 2020e), but has recently improved to endangered in response to stricter fishing quotas and the implementation of management strategies (Holdsworth, 2024). Exhibiting an epipelagic behavior, SBT are distributed throughout rich temperate waters of the Southern Hemisphere. Adults undertake extensive migrations from these feeding grounds (Block et al., 2005; Reglero et al., 2014a; Muhling et al., 2017) to reproduce in warm waters of the northeastern Indian Ocean (IO), between Java and Australia (7–20° S and 110–125° E), the only spawning ground known for SBT (Nishikawa et al., 1985; Shingu 1980; Farley and Davis, 1998; Patterson et al., 2008). Spawning occurs from September to April, with, peak larval abundances between December and March (Farley et al., 2007). Water masses in this area is characterized by the entrance of coastal, subtropical and superficial tropical water (Rochford, 1988) where the last one, from the Pacific Ocean, enters through the Indonesian Throughflow (ITF), resulting in oligotrophic conditions (Landry et al., 2022b), high temperatures and low salinity values (Domingues et al., 2007). Circulation and transport downstream of the ITF have a seasonal variations linked to the monsoon system (Clarke and Liu, 1993; Masumoto and Yamagata, 1996), and to the Leeuwin Current (LC) which plays a key role in the southern transport of larvae and juvenile SBT when they leave the spawning grounds (Nieblas et al., 2014; Hobday et al., 2015).

Larval stages represent a bottleneck to recruitment as this is the most critical period in the life of a fish, where survival is largely dependent on growth potential (Hjort et al., 1926, Houde, 1987; Anderson, 1988). The survival of fish larvae is shaped by a combination of assumptions currently encompassed in the so-called ‘Growth-Survival Paradigm’, defined by the classic assumptions of ‘*bigger is better*’ (Miller et al., 1988), ‘*stage-duration*’ (Houde, 1987, 1989), and ‘*growth-selective predation*’ (Takasuka, 2003, 2007), which relate those individuals that grow faster, or are larger at a given time, having higher probability of survival. Scombrid fishes benefit from having some of the highest larval growth rates (Morote et al., 2008), shortening their vulnerability during the early life stages. One advantage is that reaching the flexion stage faster improves swimming capacity for evading predators, optimizes feeding opportunities and may facilitate adaptability to environment fluctuations (Chick et al., 2000; Pitchford and Brindley, 2001). A higher growth potential increases the survival probability and, hence, recruitment of the species (Takasuka et al., 2003).

Growth in fishes is typically recorded in the microstructure of their otoliths, tracing the daily growth history of the earliest stages of the individuals (Campana, 1990). The applicability of otoliths for larval growth studies has been tested and analyzed in numerous studies of bluefin tuna larvae, reporting changes with larval ontogeny, nutritional condition, differences in maternal transmission and inter-annual growth variability from different spawning grounds (Jenkins and Davis, 1990; García et al., 2013; Tanaka et al., 2014; Malca et al., 2017, 2022; Hane et al., 2020; Quintanilla et al., 2023, 2024).

Sea temperature and food availability, along with maternal influence during the first stages, are the main environmental drivers that influence growth potential in bluefin tuna larvae, with higher temperature usually having a positive effect on growth (Jobling et al., 2002; Tanaka et al., 2006; Quintanilla et al., 2023, 2024) when food resources are not limited (Shropshire et al., 2022). These environmental and maternal effects are reflected in the daily width increments of otoliths (Otterlei et al., 2002) and described in numerous studies of bluefin tuna larvae (Satoh et al., 2013; Muhling et al., 2017). In contrast to other bluefin tuna species, there is relatively little recent information about SBT regarding its spawning habitat, climate characteristics, reproductive behavior and early stages, with the major study having been done in 1987 by Jenkins et al. (1990, 1991).

This study analyzes growth patterns of SBT larvae, through a detailed examination of otolith microstructure, revisiting the spawning ground at the same time of year, after three decades since the first studies. We provide a comprehensive overview of the growth of SBT larvae in the region through: i) analysis of both population-level and within-population growth in recently collected larvae; ii) comparison of current growth patterns with those reported by Jenkins et al. (1990, 1991), particularly relevant in context of climate change; and iii) an analysis of the potential implications of growth rate for larval survival.

The main objective of this work is to deepen the understanding of the early life stages of this species from a growth perspective, contributing new knowledge about larval survival and its possible influence on recruitment. This is the first population-level growth study on wild SBT larvae since those earlier investigations, allowing an update and expansion of information on the early life stages of this species.

## 2. Material and methods

### 2.1. Sample and data collection

SBT larvae were collected in 2022 during BLOOFINZ-IO cruise RR2201 on R/V *Roger Revelle*, coinciding with the spawning peak of the species (January – March) (Figure 1). At sea, larvae were identified based on morphology, meristic characteristics and pigmentation patterns following Nishikawa et al. (1985, 1987). When SBT larval patches was located (n > 5, for one side of a standard net tow), the water parcel was marked with a satellite-tracked drifter, equipped with a drogue at 15-m, and sampling was initiated at 3-6 h intervals following the drifter for 3-4 days. A four of such multi-day experiments, called cycles (Cycles 1-4, hereafter C1-C4), was conducted and during which other water-column properties and plankton rate processes were also extensively investigated (Landry et al., this issue). Larvae were also sampled at individual stations on transits between cycles and along longer, transects across the Argo Basin at the beginning and end of the cruise. SBT larvae were sampled using double oblique tows from the surface to 25 m and back for 10 min (∼2 knots) with a 90-cm Bongo net (505-μm mesh) and with a 1-m^2^ surface net (1000-μm mesh). The 90-cm Bongo net was deployed with flowmeters (General Oceanics Inc) placed at the center of each net to record volume filtered. Coupled to the Bongo 90-cm net, zooplankton prey was sampled using two 20-cm Bongo nets (200 and 50 μm mesh). The details of zooplankton sampling are described in detail by Laiz-Carrión et al. (this issue). SBT larvae were preserved by freezing (−80°C) or in 95% ethanol for their posterior analysis in Oceanographic Center of Malaga (IEO-CSIC) and University of Miami, respectively, for genetic analysis, daily growth and trophic ecology studies. Profiles of environmental variables were taken from CTD casts (Seabird SBE 9plus) using a rosette equipped with 12 Niskin bottles from the bottom to the surface. The full cruise and methodologies are described in detail by Landry et al. (this issue).

**Figure 1.**
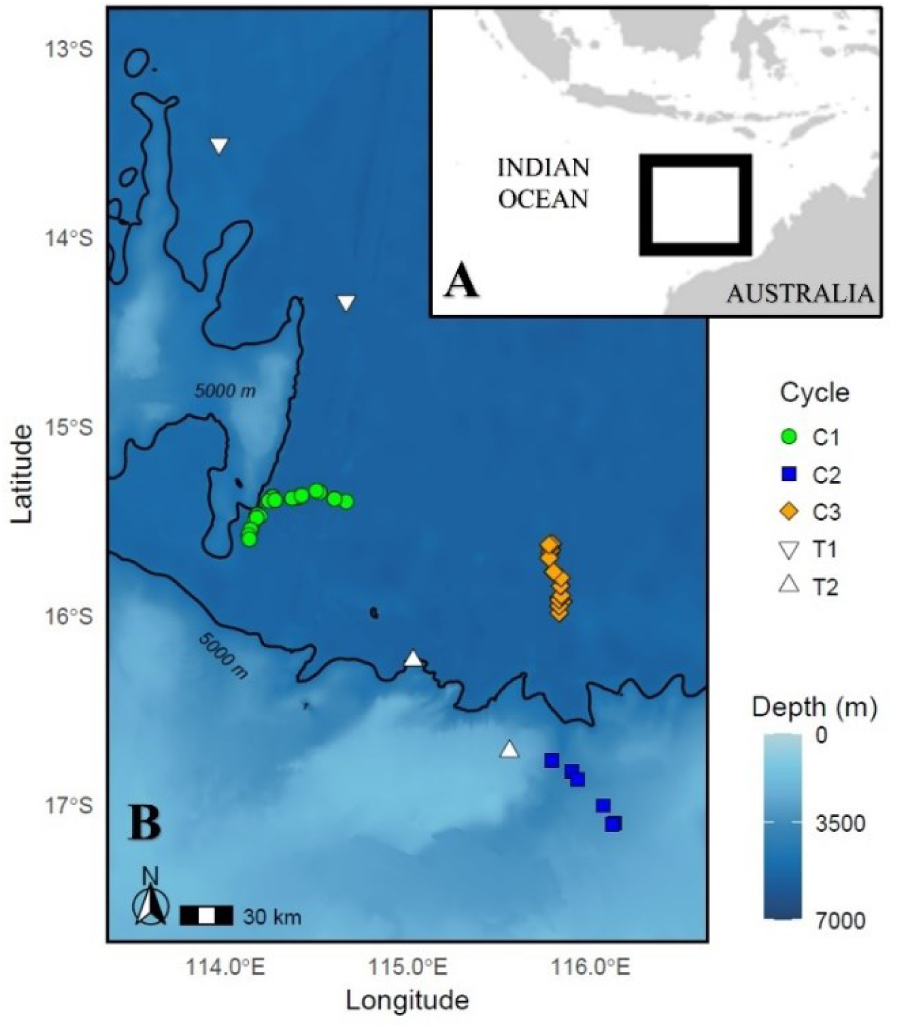
A) SBT spawning ground and B) sampling stations for larval SBT from 2022 in the eastern Indian Ocean. Locations for three cycles (C1-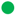 green; C2-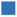 blue; C3-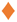 orange) and additional stations from transits (T1-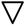, T2-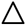) from the present study are shown.

### 2.2. Larvae and otolith processing

At the Oceanographic Center of Málaga, on-board SBT larvae identifications were reviewed by experienced larval taxonomist following the guidelines from Nishikawa et al. (1985, 1987) by experienced larval taxonomists. In addition, SBT larvae were genetically identified to confirm visual identifications. The genetic protocol and results can be consulted in Malca et al. (this issue).

Individual larvae for growth studies were night-time collected specimens, generally with empty stomachs since SBT larvae are visual predators that feed during the day (Jenkins et al., 1991; Llopiz et al., 2014; Uriarte et al., 2016). Larvae were measured for standard length (SL, mm) and body depth (BD, mm) using ImageJ software analysis (± 0.01 mm). BD was measured at an angle of 90° from the anus, following the criteria of previous studies of *T.thynnus* larvae (Catalán et al., 2011). Larval dry weight (DW, mg) was obtained with a precision microbalance (0.001 mg) after 24 h of dry-freezing following Laiz-Carrión et al. (2015). Larvae of SBT were classified based on their urostile flexion into three different development stages: pre-flexion, flexion and post-flexion (Kendall et al., 1984).

All otoliths from each larva were processed under a polarized binocular microscope following García et al. (2013). The largest, sagittal otoliths, were selected for ageing. Based on the assumption of bilateral symmetry in microstructure of left and right otoliths in other fish larvae (Campana, 1999) and tuna larvae (Jenkins and Davis, 1990; Malca et al., 2023), the best sagitta in terms of clearest daily rings and fewest artefacts were selected for aging. Otoliths were digitalized in single stacks (step size = 1 μm), to a depth that depended on otolith size (Quintanilla et al., 2023, 2024), using the optical microscope Leica CTR6 LED at 1000x magnification and the image software analysis Leica Application Suite X LAS X 2.0.0. We measured otolith biometrics of radius (RD, μm) and daily increments width (IW, μm), the latter used to calculate the mean increment width (MIW, μm). Daily increments (AGE, days post hatching-dph) were counted and estimated following the reading criteria for Atlantic Bluefin Tuna larvae (ABFT) detailed in Malca et al. (2023), and carried out in previous studies in tuna larvae (García et al., 2013; Quintanilla et al., 2023, 2024).

### 2.3. Growth Analysis

SBT larval growth was determined from the somatic and otolith variables using a linear regression model (*y=a+b*x*) with AGE as the independent variable. Next, the growth variables were examined against BD to test for any correlation between somatic and biometric otolith variables. Lastly, SBT larvae were also examined at the intra-population level and divided in two groups: those with optimal (OPT) or deficient (DEF) growth with respect the whole population was selected, following Quintanilla et al. (2015), and utilized for bluefin tuna larvae (Malca et al., 2023; Quintanilla et al., 2023, 2024). The groups were divided based on positive and negative residuals of SL and DW versus AGE, with OPT larvae having positive residuals from both regressions and DEF having negative residuals. An analysis of covariance (ANCOVA) was performed to compare the somatic and otolith variables between OPT and DEF growers as factors, and AGE as covariable. The flexion stage is a critical development stage because it coincides with the beginning of exogenous feeding, which involves a significant change in larval growth (Miyashita et al., 2001; Uriarte et al., 2016). In the entire population, as well as in OPT and DEF groups, a binomial model was applied between pre-flexion and flexion stage to model the probability of an individual of being in either the pre-flexion or flexion in relation of standard length, age and dried weight, as continuous variables. Next, we used this approach to estimate the point where 50% of the larval population reached the flexion stage for the standard length (SL_F50_), age (AGE_F50_) and dried weight (DW_F50_).

### 2.4. Retrospective analysis

Our growth pattern was compared with Jenkins and Davis (1990) with an ANCOVA-base retrospective approach using AGE as covariate. The theorical SL of the historical study was calculated based on our estimated AGE, using the linear equation from Jenkins and Davis. (1990). To make both growth studies comparable, a common range of SL was selected, from a previous transformation to SL in ethanol (SL_ETHANOL_) by the ethanol shrinkage curve across (Eq. 1). This curve was based on ∼100 SBT larvae that were initially measured fresh at sea for SL, and later re-measured in the laboratory after preservation in 95% ethanol (N = 87). This methodology follows a similar approach used by Malca et al. (2017, 2022) for ABFT larvae.

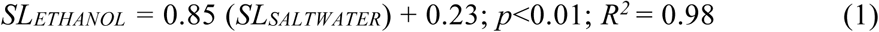

### 2.5. Survival insights

A field-based survival study for SBT larvae was conducted sampling the same water mass on two different occasions during the cycle studies. After finalizing C1, the water mass was marked with a drogued surface drifter to be resampled later during C3 to assess the growth potential and survival of SBT larvae. Larvae of SBT were collected in both cycles (C1, C3) and a survival experiment was carried out by back-calculating day of hatching for each larva of both cycles, based on assigned AGE. Thus, we assumed that larvae collected during both cycles came from the same water mass, and those hatched during the same week belonged to the same cohort, which allows us to distinguish between the initial population (C1) and the older larvae that survived and were sampled later (C3). Larvae that matched these criteria were selected and their IW were compared as a proxy of potential growth. C1 was divided in OPT and DEF groups, to compare the IW of the otolith from C1 and OPT with survived larvae from C3. A decreasing percentage of larvae from C1 with higher or same mean value of IW than C3 were estimated to elucidate the number of potential survived larvae with similar growth pattern as C3. In a similar approach to exploring growth patterns, the OPT group from C1 was divided in four subgroups based on the percentage of the fastest-growing larvae: 100%, 75%, 50%, and 25%. The mean increment width (MIW) of each category was then examined and compared to that of the C3 group using the non-parametric Mann-Whitney U test to assess differences between the groups.

### 2.6 Environmental variables

The environmental data used, collected within the upper 25 meters, were obtained from the casts closest to the larval stations. Temperature (°C), salinity (psu), mixed-layer depth (m), and the biomass of micro- and mesozooplankton (mg/m³) were analyzed with a Mann-Whitney U tests both between OPT and DEF stations within the larval population, and between stations of groups from survival insights, in order to characterize any environmental differences that may explain larval growth variability.

Statistical analyses were carried out using R version 4.2.1 (R Core Team 23-6-22 ucrt) through the integrated development environment RStudio, with α = 0.05. When necessary, variables were log-transformed to obtain the linearity and variance homogeneity assumptions (Sokal and Rohlf, 1971).

## 3. Results

### 3.1 Larval growth

SBT larvae were collected from three cycles (C1=73; C2=8; C3=94) and two transits (T1=4; T2=6), with the majority from C1 and C3 (Fig. 1). A total of 185 night-time SBT larvae from 44 stations were aged for growth study. The freshly SL and AGE range for SBT larvae of the population was ranged from 2.5 to 11 mm (Fig. 2A) and 4 to 23 dph (Fig. 2B), respectively.

**Figure 2.**
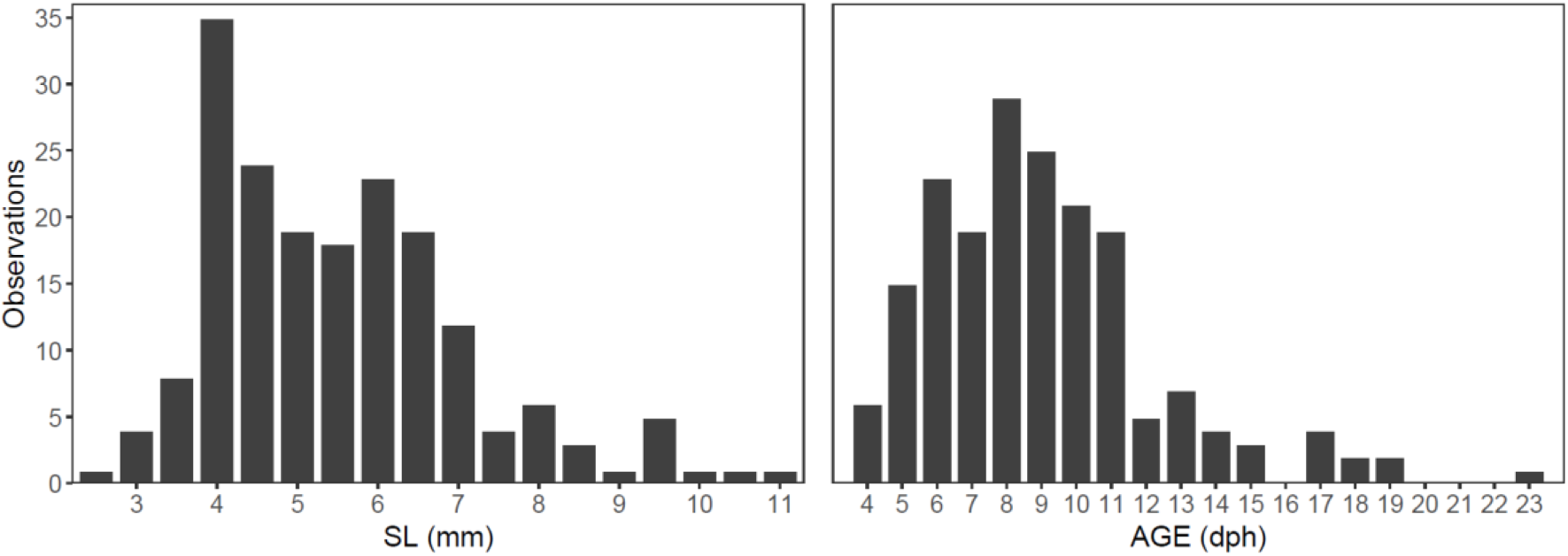
Histograms for SBT larvae (N=185) for (A) standard length (SL, mm) and (B) age days post hatch (dph) of frequencies for (A) standard length and (B) age in SBT larvae.

The means and standard deviations (Mean ± Sd) were calculated for somatic and otolith biometric variables for SBT larvae and summarized in Table 1. Both somatic and otolith biometric variables showed a linear positive relationship and statistical significance with AGE. The growth rate was 0.46 mm d^-1^ (Table 1; Fig. 3). BD displayed a strong linear relationship with the rest of larval somatic (SL, DW) and otolith (RD, MIW) variables (Table 2; Fig. 4).

**Figure 3.**
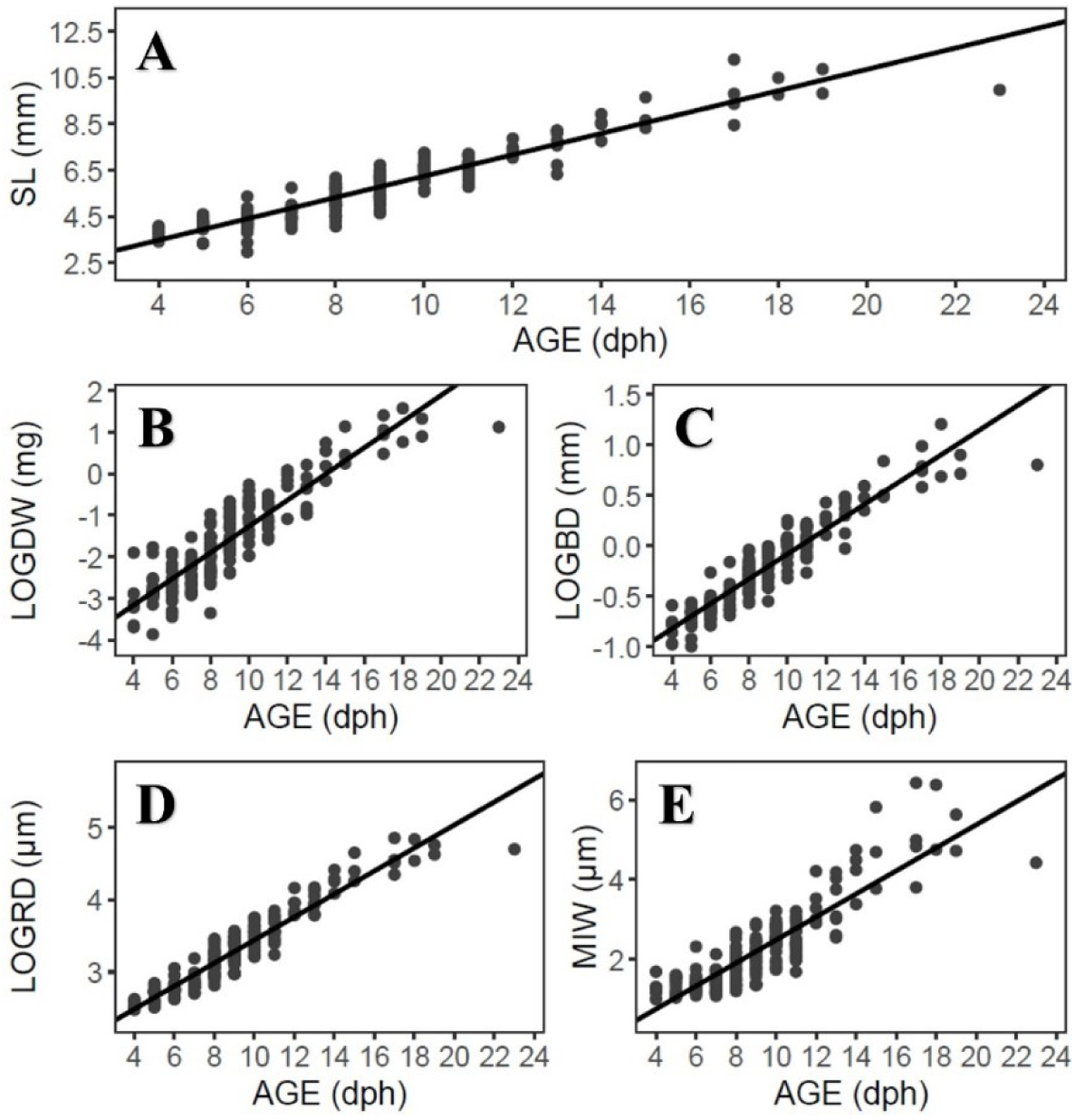
Least square regressions for somatic (A, B, C) and otolith variables (D, E) of SBT larvae. The (A) standard length, (B) dry weight, (C) body depth, (D) otolith radius and (E) mean increment width versus AGE are shown for the total population. Some of the variables were log-transformed (LOG).

**Figure 4.**
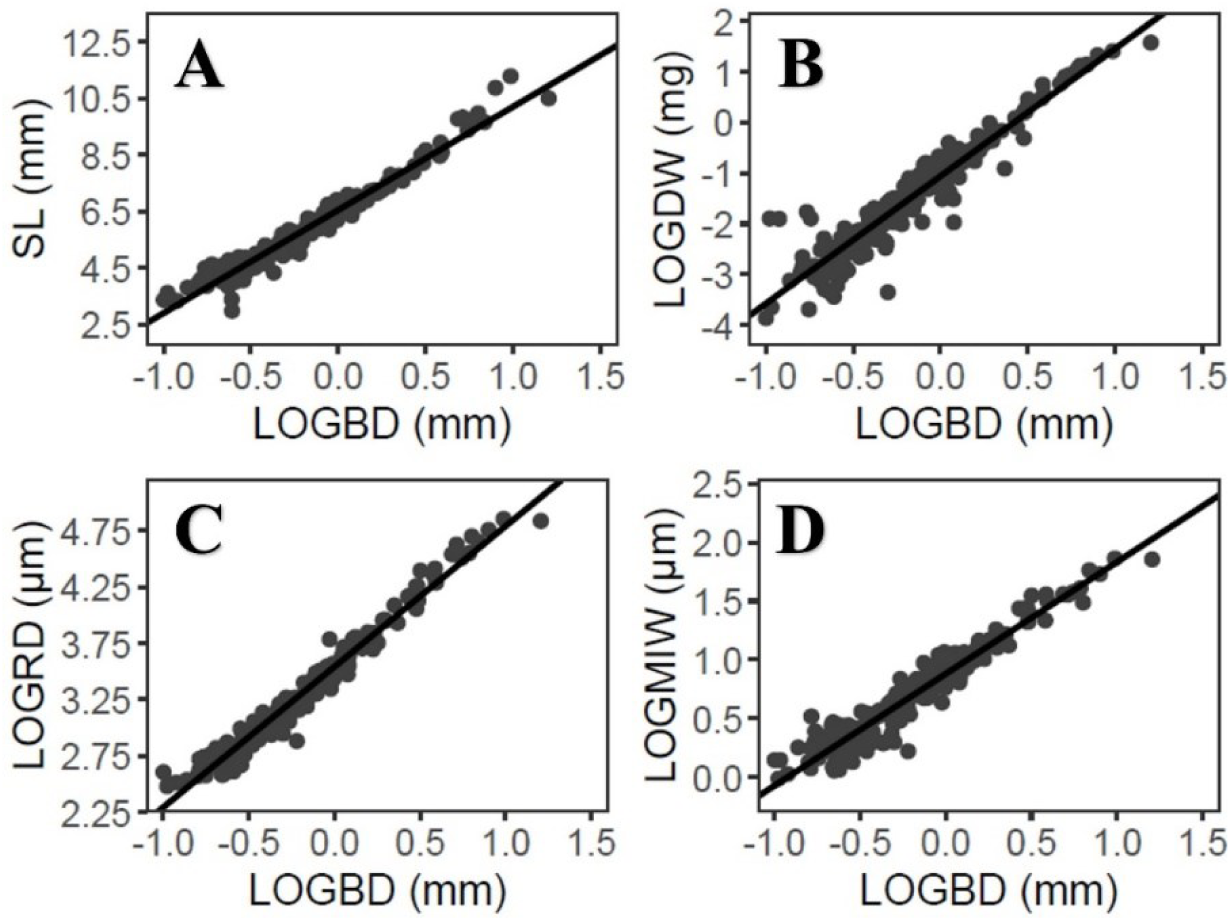
Least square regressions for somatic (A, B) and otolith variables (C, D) of SBT larvae. The (A) standard length, (B) dry weight, (C) otolith radius and (D) mean increment width versus body depth (BD) are shown for the total population. Some of the variables were log-transformed (LOG).

**Table 1.**
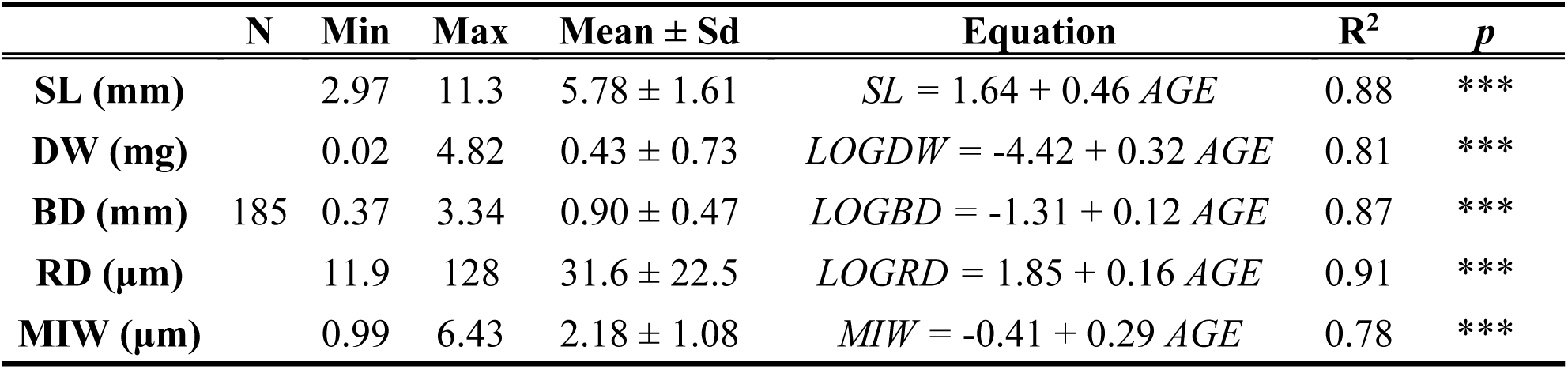
Summaries of somatic (standard length - SL,mm; dry weight - DW, mg; body depth – BD, mm) and otolith biometric variables (radius - RD, μm; mean incement width - MIW, μm) for SBT population. Number of individuals (N), minimum (Min) and maximum (Max) values together with mean and standard deviation (Mean ± Sd) are showed. The linear regression equations (Equation), R^2^ and *p*-value (*p*) with AGE (dph) are represented. *** p-value < 0,001. Some of variables were log-transformed (*LOG*).

| | N | Min | Max | Mean ± Sd | Equation | R<sup>2</sup> | *p* |
| --- | --- | --- | --- | --- | --- | --- | --- |
| **SL (mm)** | | 2.97 | 11.3 | 5.78 ± 1.61 | $SL = 1.64 + 0.46 AGE$ | 0.88 | *** |
| **DW (mg)** | | 0.02 | 4.82 | 0.43 ± 0.73 | $LOGDW = -4.42 + 0.32 AGE$ | 0.81 | *** |
| **BD (mm)** | 185 | 0.37 | 3.34 | 0.90 ± 0.47 | $LOGBD = -1.31 + 0.12 AGE$ | 0.87 | *** |
| **RD (μm)** | | 11.9 | 128 | 31.6 ± 22.5 | $LOGRD = 1.85 + 0.16 AGE$ | 0.91 | *** |
| **MIW (μm)** | | 0.99 | 6.43 | 2.18 ± 1.08 | $MIW = -0.41 + 0.29 AGE$ | 0.78 | *** |

**Table 2.**
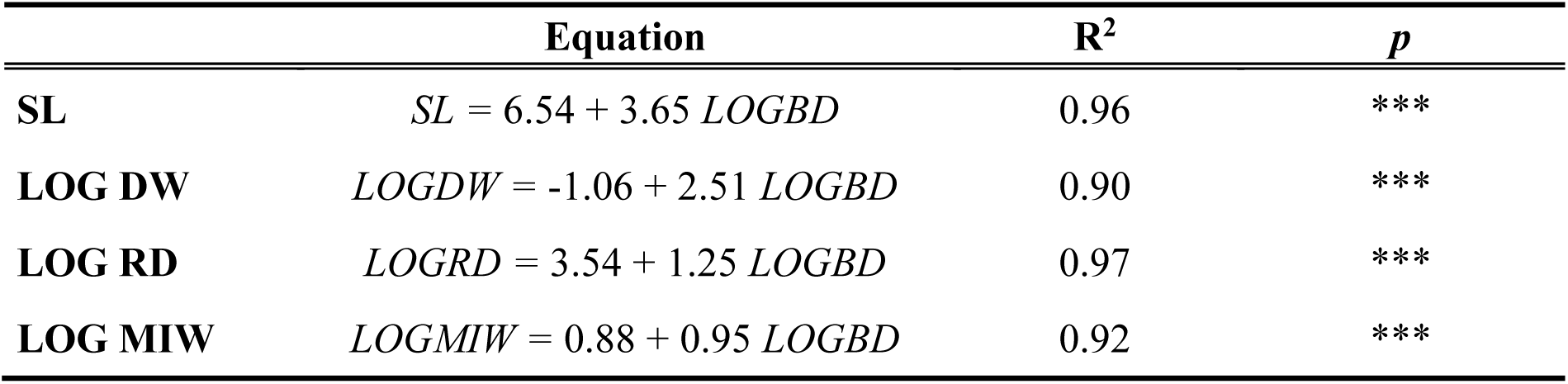
Linear regression equations (Equation), R^2^ and *p*-value (*p*) with body depth (BD, mm) are represented. *** p-value < 0,001. Some of variables were log-transformed (*LOG*).

The larval population was equally distributed between OPT and DEF groups, which each representing 38% of the total individuals. In the OPT group, individuals showed higher values, compared to DEF group (ANCOVA, *p* < 0.001), of SL (6.43 ± 1.30 vs 5.21 ± 1.67) and DW (0.55 ± 0.64 vs 0.35 ± 0.74), as well as greater BD (1.04 ± 0.40 vs 0.77 ± 0.46 mm), RD (36.80 ± 19.80 vs 27.70 ± 24 μm), and MIW (2.64 ± 0.97 vs 1.81 ± 1.02).

The flexion stage was characterized by estimation of SL, DW and AGE of flexion for total population, OPT and DEF groups. Our model estimated that 50% of the SBT population reached the larval flexion at SL_F50_ 5.4 mm, a DW_F50_ 0.2 mg and AGE_F50_ 8.5 days. Regarding the OPT and DEF groups, higher values of SL_F50_ and DW_F50_ were observed in the OPT group, but similar values of AGE_F50_ (differing only by ∼1 day) (Table 3; Fig. 5).

**Figure 5.**
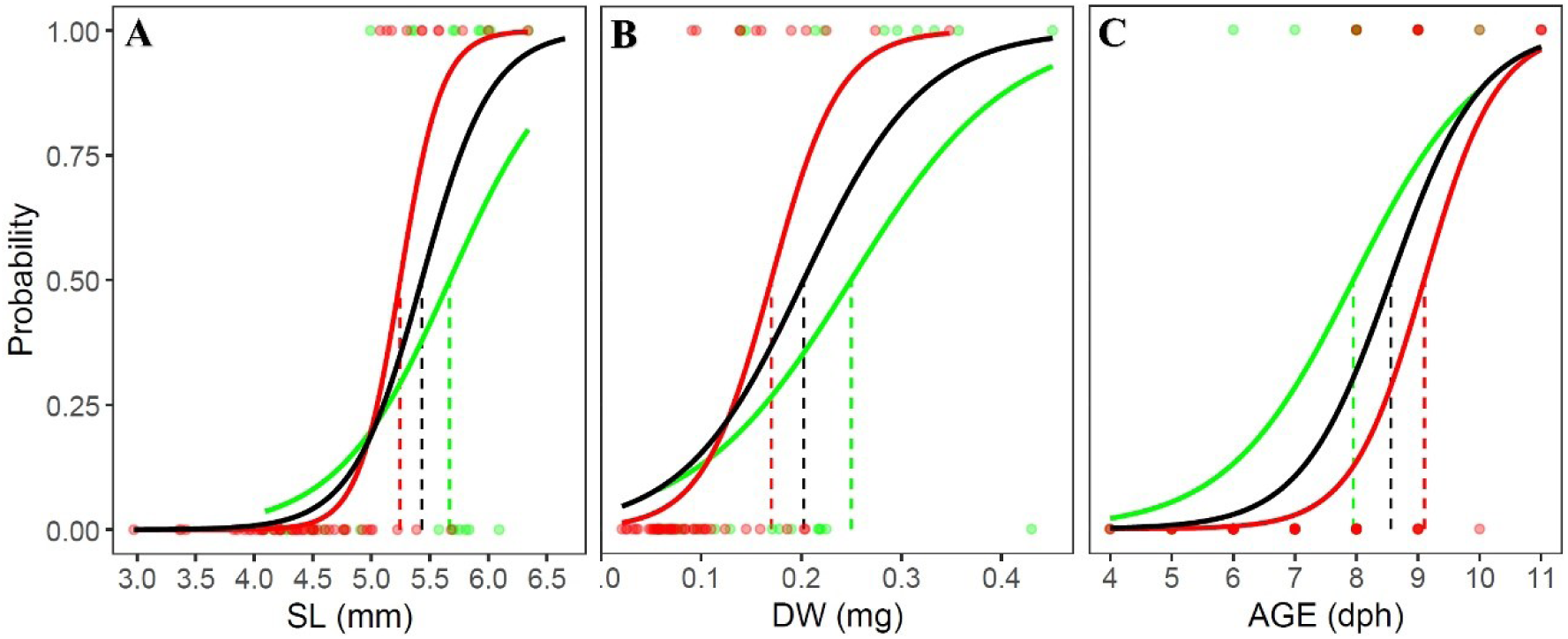
(A) Standard length (SL_F50_), (B) dry weight (DW_F50_) and (C) age (AGE_F50_) of flexion for SBT larval population (Total – **black**) and optimal (OPT – **green**)) and deficient (DEF – **red**) groups.

**Table 3.** Results of larval SBT standard length (SL_F50_), dry weight (DW_F50_), and age (AGE_F50_) of flexion for the total population, optimal (OPT) and deficient (DEF) groups, obtained from a conducted model between pre-flexion (Pre) and flexion (Flex) stages. Observations of Pre and Flex, and SL_F50_, DW_F50_, and AGE_F50_, Confident Interval (C.I) and R^2^ are showed.

| | Pre | Flex | $SL_{F50}$ | C.I | $R^2$ | $DW_{F50}$ | C.I | $R^2$ | $AGE_{F50}$ | C.I | $R^2$ |
| --- | --- | --- | --- | --- | --- | --- | --- | --- | --- | --- | --- |
| **Total** | 85 | 30 | 5.43 | 5.2 - 5.6 | 0.84 | 0.2 | 0.1 - 0.2 | 0.74 | 8.50 | 7.9 - 8.6 | 0.78 |
| **OPT** | 17 | 10 | 5.67 | 5.2 - 5.8 | 0.71 | 0.25 | 0.2 - 0.3 | 0.71 | 7.90 | 6.7 - 8.2 | 0.71 |
| **DEF** | 47 | 11 | 5.25 | 5.0 - 5.5 | 0.86 | 0.17 | 0.1 - 0.2 | 0.75 | 9.00 | 8.5 - 9.6 | 0.79 |

Environmental variables influencing the growth of the SBT larval population were characterized at each sampling station. SBT larvae were found in waters with an average temperature of 29.0 ± 0.4 °C and a mean salinity of 34.6 ± 0.14 psu. In reference to stations of OPT (N = 32) and DEF (N = 32) groups, no significant differences in temperature (29.0 ± 0.41 vs 28.9 ± 0.42), salinity (34.6 ± 0.12 vs 34.6 ± 0.13), mixed layer (16.5 ± 15.5 vs 19.0 ± 15.9), biomass of microzooplankton (1.08 ± 0.56 vs 1.33 ± 1.04) and mesozooplankton (6.30 ± 3.97 vs 5.94 ± 3.41) were found (Mann-Whitney U test, p > 0.05).

### 3.2. Retrospective analysis

The shrinkage correction for SBT larvae measured in saltwater for 95% ethanol was 10.77% ± 3.72 (Mean ± SD), obtained from 87 saltwater larvae with standard lengths between 2.68 and 11.80 mm. Upon, applying the shrinkage correction the SL_ETHANOL_ ranged from 2.49 to 10.56 mm, with a mean value of 5.43 ± 1.78. For the retrospective analysis, SL_ETHANOL_ ranged similarly from 3 to 10 mm following Jenkins and Davis (1990), but significant differences were found with higher SL_ETHANOL_ values in SBT larvae from 2022 (ANCOVA; F_1,366_ = 66.9; *p* < 0.001). In terms of growth rates, SBT from 2022 exhibited higher growth rate values (0.38 mm d⁻¹) than those from 1987 (0.33 mm d⁻¹).

### 3.3 Survival insights

Survival experiment was performed from back-calculation analysis, showing an initial (C1; N=69) and a survived cohort (C3; N=11) that were born in the same week, between 25 January and 2 February.

The IW of otoliths from the initial cohort had narrower increments when compared to the same early increments of the survivors. Initially, during the first two days post hatch, less than 40% of larvae had similar or larger IW than those of the survivors; however, this proportion declined over time, reaching less than 2% (∼1.47%) by increment 7 (Fig. 6A). Regarding the OPT groups, from the onset, only 35 – 49% of initial larvae had similar or larger IW than survivors, with a steady decreasing trend until reaching increment 9, where only the ∼4-5-7-14% of larvae from OPT subgroups 100-75-50-25%, respectively, had similar or larger IW than survivors (Fig. 6B).

**Figure 6.**
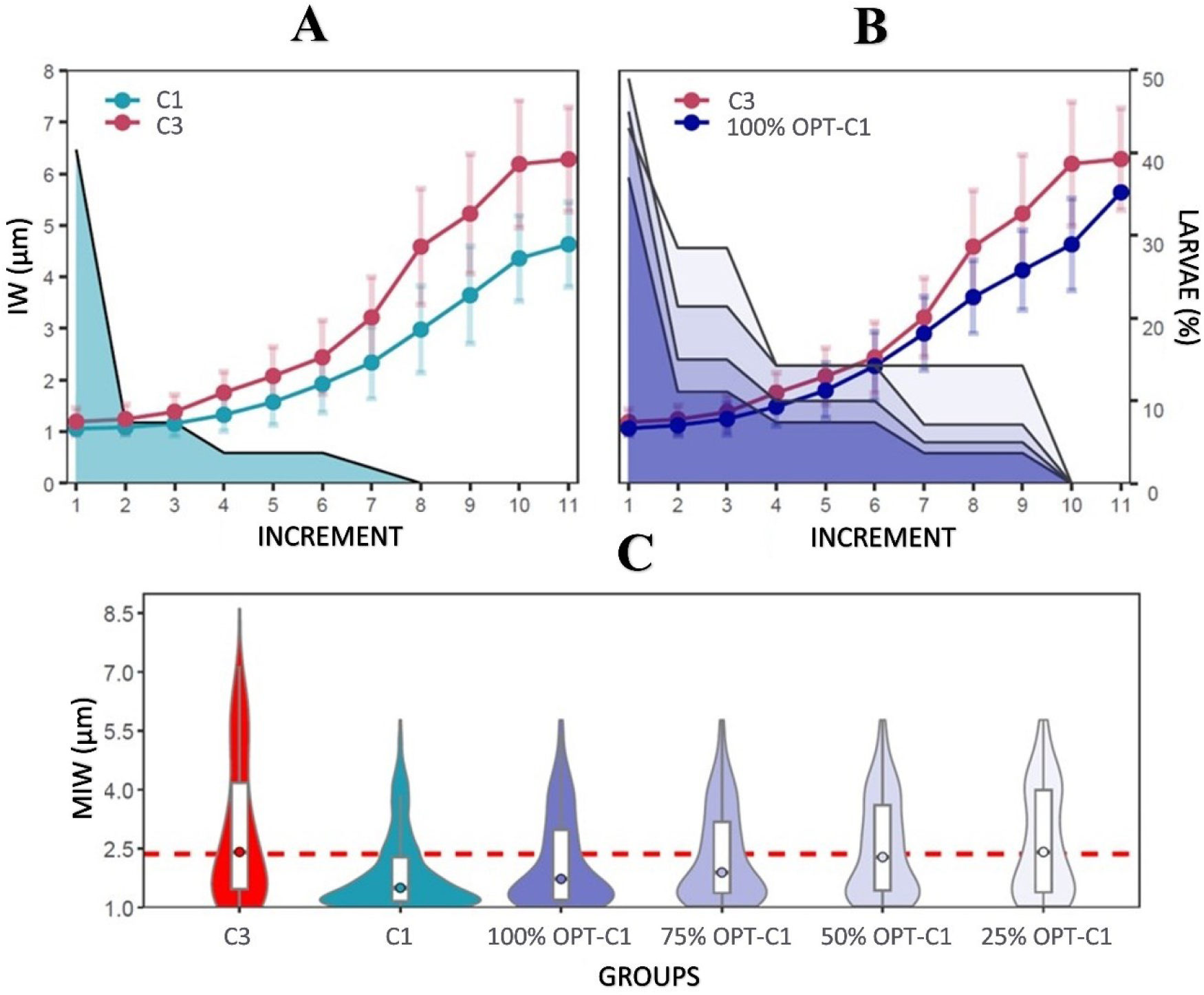
(A) Mean values for daily increments width (IW, μm) for initial population (C1, blue) and survivors (C3, red). Shaded blue area represents the percentage of larvae from C1 with similar or higher IW than C3 at each increment. (B) Mean values for daily increments width (IW, μm) for C3 (red) and 100% OPT-C1 (dark blue). Shaded blue gradient represents the OPT subgroups (100% OPT-C1, 75% OPT-C1, 50% OPT-C1, 25% OPT-C1) with larvae with similar or higher IW than C3 at each increment. (C) Boxplot and violin plot for mean increment width (MIW) for survivors (C3), the initial population (C1) and their different OPT groups (100% OPT-C1, 75% OPT-C1, 50% OPT-C1, 25% OPT-C1). The points represent the median value and the red dashed horizontal line is the median IW for the C3 group.

The MIW from the initial cohort were significantly lower than for survivors (Mann-Whitney U test, *p*<0.05). Although the OPT subgroups (100%, 75%, 50%, 25%) showed progressively higher MIW values, they still did not reach MIW equal to the survived larvae, except for the 50% and 25% OPT whose median values were similar to the survivors (Mann-Whitney U test, *p*>0.05) (Fig. 6C).

No significant differences in environmental variables (Mann-Whitney test, p > 0.05) were observed among the sampling stations corresponding to the initial cohort (C1) and those with surviving larvae (C3).

## 4. Discussion

Understanding the recruitment dynamics of SBT is crucial for ensuring long-term sustainable resource exploitation and managements of this endangered species (Collette et al., 2021). Despite signs of stock recovery (Bravington et al., 2016; Anon, 2023), consistent larval and recruitment monitoring remains scarce. This lack of routine larval sampling presents a critical knowledge gap in evaluating SBT recruitment variability and early life survival mechanisms. As a long-lived species characterized by slow growth and late maturity (Gunn et al., 2008), SBT larvae are in sharp contrast sharply with remarkably rapid development and accelerated growth (Jenkins and Davis., 1990; Gunn et al., 2008; Farley et al., 2014). This study contributes to the BLOOFINZ-IO program, representing a significant step forward in updating larval growth parameters of SBT, based on the most recent collections since the historical 1987 survey. By revisiting otolith-based daily growth, we gain a contemporary perspective on the developmental potential of SBT larvae under present-day environmental conditions.

### 4.1.1. Population growth

SBT larvae in our study exhibited a growth rate of 0.46 mm d⁻¹, encompassing a representative portion of the spawning period with conditions averaging 29 °C and 34.6 psu for temperature and salinity, respectively, in the upper 25 m of the water column. Temperature is widely considered as the main factor driving variability in larval development and growth rates (Robert et al., 2023), enhancing both processes during their early life (Pittman et al., 2013). Larval habitats of *Thunnus* species share common characteristics: they are oligotrophic zones with warm water temperatures and low primary production, conditions that increase larval survival (Teo et al., 2007; Alemany et al., 2010; Muhling et al., 2017). However, variability has been reported for larval growth rates between different bluefin spawning areas. The growth rate reported here aligns with upper estimates from the Gulf of Mexico (0.40–0.46 mm d⁻¹) and exceeds those from the Mediterranean (Balearic Sea, 0.26– 0.38 mm d⁻¹), both at slightly lower temperatures, 27°C and 23.8-26 °C respectively (García et al., 2013; Laiz-Carrión et al., 2015; Muhling et al., 2017; Malca et al., 2017, 2022). Similar growth rates have been reported for Pacific bluefin tuna (PBT) and Yellowfin tuna (YFT) larvae, though generally at cooler temperatures (<29 °C) (Foreman, 1980; Laurs et al., 1985; Tanaka et al., 2006; Satoh et al., 2008). In this context, SBT larvae show comparable growth rates, within a warmer thermal regime for the *Thunnus* species’ spawning grounds. Although high temperatures can favor larval growth when prey are abundant, they may become a limiting factor under suboptimal feeding conditions, acting as a double-edged sword depending on the ecological scenario (Reglero et al., 2025). In our study, SBT larvae were found at relatively high average temperatures compared to those reported for other *Thunnus* species. Nevertheless, the observed growth rates suggest that food availability was sufficient to support accelerated growth under these warm conditions (Shropshire et al., 2022; Swalethorp et al., *this issue*).

Larval growth was also assessed by dry weight and otolith microstructure examination, as has been done in other bluefin tuna larvae studies (Itoh et al., 2000; Quintanilla et al., 2024). However, changes in BD during SBT early life stages were document for the first time in this study. We observed a strong correlation with BD and other somatic and biometric otolith variables, supporting its potential use as a proxy for larval condition (Fig. 4; Table 2). Previous studies on bluefin tuna larvae in the Gulf of Mexico have shown a strong association between BD and the most recently formed otolith increments (recent growth); moreover, while no interannual differences were observed, heavier larvae exhibited larger BD (Malca et al., 2022). Based on our results, the strong relationship between dry weight and body depth suggested the potential for using BD as a proxy for dry weight in SBT larvae, and, given the findings of Gleiber et al. (2020) and Malca et al. (2022), possibly for other *Thunnus* species as well. Importantly, BD is not only a proxy for larval condition, but also an index of musculature development. When measured from the outer margin of the ventral-most myomere to the dorsal-most myomere, BD reflects what is sometimes referred to as “muscular height.” Thus, larvae with greater BD have more muscle mass and are, by extension, in better physical condition. This makes BD a particularly informative metric for assessing larval fitness, especially in contexts where direct weight measurements are compromised, such as in ethanol-preserved samples. This approach would be especially useful for estimating DW in such specimens, also considering the shrinkage size model proposed in this study. A strong correlation with the standard length (SL), radius (RD) and mean increment width (MIW) was also observed further support its utility as a proxy metric, particularly valuable for preserved larvae where shrinkage may compromise other morphometrics variables or damaged fish larvae during sampling, resulting in deterioration or partial fragmentation (Paine et al., 2007). The significant relationship between SL and BD could allow for the estimation of larval SL when it cannot be directly measured.

The flexion stage is particularly important, as it marks a key ontogenetic transition, between pre-flexion and post-flexion stages, characterized by concurrent morphological, physiological, and trophic changes (Anderson et al., 1988; Tanaka et al., 2006; Morote et al., 2008; Uriarte et al., 2016). For the first time for SBT, the point at which 50% of larvae reached flexion was estimated at size of approximately 5.43 mm (SL_F50_), weight of 0.20 mg (DW_F50_), and at 8-9 dph (AGE_F50_). Some previous aquaculture studies on bluefin tuna larvae reported flexion at ∼6.7 mm and ∼8 dph at 28 °C (Blanco et al., 2019), while at lower temperatures (∼24 °C), flexion occurred at smaller sizes and several days later, between 5.5–6.5 mm and around 15 dph (Uriarte et al., 2016). In wild bluefin tuna larvae, from different spawning grounds, flexion has been observed between 6–7 mm at similar lower temperatures (Malca et al., 2023) and for PBT the flexion has been described at 5– 5.7 mm at 24 – 27°C (Miyashita et al., 2001). In this context, the flexion SL observed in our study aligns within the range reported for other *Thunnus* species, with slight variations likely due to species-specific traits or environmental factors, particularly temperature. In the IO, SBT larvae were exposed to ∼29 °C during early development, which may have accelerated their ontogenetic development, as higher temperatures are known to enhance metabolic rates (Pittman et al., 2013).

### 4.1.2. Intra-population growth

At the intra-population level, distinct growth trajectories were evident between the OPT (optimal) and DEF (deficient) groups in our study. However, no significant differences in environmental variables were found among the stations where fast- and slow-growing larvae were collected. Larvae from the OPT group were significantly larger, heavier and had otoliths with wider radius and increment width, indicating a higher growth potential. With regard to the timing of notochord flexion, both groups reached flexion at roughly the same age (a ∼1-day difference is within an expect-level of subjectivity (Cardinale and Arrhenius, 2004)), but OPT group did so at greater length, weight (Fig. 5A, 5B, 5C) and body depth, suggesting higher growth rates. This observation is relevant, as length and weight are commonly used as proxies for larval condition, with higher values typically associated with improved condition and survival (Garrido et al., 2015). The OPT group would therefore reflect a better physical condition at the time of flexion.

Despite our results suggesting that flexion may be a programmed process, the physical condition at which larvae reach flexion can still be influenced by environmental variables, particularly temperature (Tsuda et al., 2012), but also food availability (Takahashi et al., 2004; Laiz-Carrión et al., 2019, this issue; Shropshire et al., 2022) and maternal inheritance (Quintanilla et al., 2023; 2024, this issue), especially during the earliest developmental stages. Maternal effects are especially important during the earliest stages, as both egg quality have a direct influence larval survival and initial growth (Ohshimo et al., 2018a; Hiraoka et al., 2019). These effects gradually decline as larval development advances, and the exogenus feeding begins (Green and McCormick, 2005; Shiroza et al., 2022). In scombrids, the timing of first feeding onset and prey availability is considered a particularly critical point in larval development, as it directly influences future feeding success and larval growth (Pepin et al., 2015).

The higher potential growth and better larval condition observed at the flexion stage in the OPT group might be partially explained by improved maternal condition or differences in maternal behavior (Sakai and Harada, 2001), as well as by variations in larval trophic behavior, as previously reported at the population level and among fast- and slow-growing pre-flexion ABFT larvae (Quintanilla et al., 2023, 2024, this issue). These findings highlight the importance of larval physical condition at the flexion stage, as this is a phase of increased energy demand where a good condition can enhance survival by improving feeding capacity, predator evasion, and resilience to short-term starvation (Chick et al., 2000).

### 4.2. Retrospective comparison

Since the pioneering studies by Jenkins and Davis et al. (1990) and Jenkins et al. (1991) published over three decades ago, no further studies had been conducted on the larval growth dynamics of SBT within their only known spawning ground. SBT larvae from 2022 had a higher larval growth potential (0.38 mm d^-1^) compared to 1987 (0.33 mm d^-1^), possibly linked to differences in environmental conditions, including an increase of ∼2 °C in sea surface temperature. Considering the key role of temperature in larval development of tunas (García et al., 2013), this warming may partially explain the higher growth rates observed, assuming sufficient food availability (Gleiber et al., 2020). However, the increased growth observed in SBT larvae in recent years may not be solely attributed to higher temperatures, but also to food availability (Govoni et al., 1985). In 2022, zooplankton diversity differed and trophic resources were higher, with a total abundance of 2.7-fold higher than in 1987, which may have resulted from variations in prey nutritional quality (Davies et al., this issue) or from shifts in larval active prey trophic behavior (Laiz-Carrión et al., this issue; Swalethorph et al., this issue), playing critical roles in supporting the higher metabolic demands associated with elevated growth rates.

Nevertheless, when food resources are insufficient or thermal threshold are exceeded, elevated temperatures can negatively impact larval development (Tanaka et al., 2006; Pittman et al., 2013; Satoh et al., 2013). A broad upper thermal limit has been identified for various tuna larvae species, typically between 30-32 °C, beyond which, larval survival and development decline markedly with increased rates of malformations and mortality (Gleiber et al., 2020; Ortega et al., 2024). SBT larvae were found in water temperatures at 29 °C, substantially higher than the 27.2–27.7 °C reported in 1987, approaching the upper limit considered optimal for larval development for bluefin tunas (Ortega et al., 2024; Reglero et al., 2025). This suggests that although current conditions may still support viable larval development at the spawning grounds, even slight increases in temperature could push conditions beyond the larvae’s functional thermal range, compromising survival and recruitment success. The increase in water temperature can affect pelagic trophic dynamics (Houde, 1987; Buckley et al., 2004), with potential negative impacts on species whose early life stages occur in tropical, food-limited conditions (Llopiz and Hobday, 2015). The reproductive behavior of adults could also be affected, as rising temperatures in the spawning grounds may alter reproductive patterns if optimal thermal conditions are no longer met (Schaefer, 2001; Locarnini et al., 2010; Rooker et al., 2014). These thermal limits are particularly critical in the current framework of global climate change, which is driving rising temperatures across all oceans, especially in the IO, where warming trends are accelerating among the fastest-warming oceans on the planet (Fox-Kemper et al., 2021; Roxy et al., 2024).

### 4.3. Survival insights

Notable differences between the initial cohort (C1) and the surviving cohort (C3) were found. Using otolith increment width (IW) as a reliable indicator, our results showed that larval survival of SBT was strongly associated with growth. In this sense, wider increments and larger MIW observed for the surviving cohort (C3) (Fig. 6A), supporting the “growth-selective survival” hypothesis, wherein larvae with larger increments exhibit higher somatic growth rates and, consequently, a greater likelihood of survival (Allain et al., 2003). This pattern has been documented also in BFT, demonstrating a close relationship between otolith development and somatic growth throughout the larval stage (Jenkins et al., 1991; Malca et al., 2017; Watai et al., 2017; Quintanilla et al., 2023).

During the first days of larval life, differences in increment widths were already evident. From the onset of ontogeny, fewer than half of the initial larvae (C1) exhibited increment sizes comparable to those of the surviving larvae (Fig. 6A), indicating a clear selection for growth potential immediately post hatch. Larval growth is highly variable and can depend on both extrinsic and intrinsic factors. Temperature plays a crucial role, as it affects yolk utilization efficiency and can ultimately influence larval size at hatching (Pepin et al., 1997; Peck et al., 2012). However, in our study, no differences in mean temperature were observed between stations, suggesting that other factors may be primarily responsible for the observed differences in increment widths for initial and survived cohort. Especially in the pre-flexion stage, maternal influence has been shown to play a key role in the survival of ABFT larvae (Quintanilla et al., 2023; 2024) and SBT larvae (Quintanilla et al., this issue). Based on this, the surviving larvae may have had, from the very beginning of their life, a higher growth potential, reflected in enhanced pre-flexion growth, directly inherited from their mothers (Fennie et al., 2024; Walsh et al., 2024).

As larval development progresses, the difference in increment width remained consistent up to approximately day 7, at which point, the differences between initial and survivor cohort became more pronounced (Fig. 6A). This shift coincides roughly with the onset of flexion (Fig. 5C) and was accompanied by dietary shifts (Shiroza et al., 2022; Swalethorp et al., this issue). Fewer than 2% of larvae attained IW comparable to the survivors by that day, beyond which no larvae achieved increment sizes similar to those of the cohort survivors. This low larval survival may be attributed to suboptimal environmental conditions during the early developmental stages. Just prior to Cycle 1, the water column had experienced intense mixing by a tropical storm (Landry et al., this issue b), which may have dispersed both prey and larvae, reducing prey–larva overlap. In contrast, stratified water conditions have been shown to favor bluefin tuna larvae by concentrating both prey and larvae in surface layers (Röpke, 1993; Satoh et al., 2013), facilitating more efficient foraging. This hypothesis is further supported by differences in plankton composition (Davies et al., this issue), larval densities (Malca et al., this issue) and stomach content analysis (Swalethorp et al., this issue) between Cycles 1 and subsequent cycles more stratified. The mixing event of the water column during C1 may have imposed trophic limitations on larvae by reducing prey availability and increasing starvation risk, and in consequence the surviving of larvae. Larvae from the initial cohort likely had lower growth potential from hatching compared to the survivors, which may have compromised their survival under such environmental constraints. In contrast, survivor larvae have a higher growth potential from hatching and probably a larger at hatch. The bigger the larvae are when it hatches the more likely it is to grow fast and survive, and continue to grow fast later in life (Garrido et al., 2015; Fennie et al., 2024). Rapid early growth at the beginning of its life may have been better able to withstand starvation and capture larger, more nutritious prey, thus enhancing their chances of survival. Our findings highlight the critical role of maternal inheritance in early-life growth and survival of SBT, especially under unfavorable environmental conditions (Meekan and Fortier, 1996; Grønkjær and Schytte, 1999; Vigliola et al., 2002; Raventos et al., 2005). The early rapid growers tend to develop superior swimming and predatory abilities, enabling predator evasion, selective feeding, more efficient energy utilization and, in consequences, major survival probability in flexion and postflexion stages (Laiz-Carrión et al., this issue). This supports the idea of a selective process favoring individuals with higher growth rates from the earliest larval stages (Takasuka et al., 2017; Quintanilla et al., in review), underscoring the survival advantage conferred by higher growth potential and better resource exploitation.

Regarding the fast-growing larval group (OPT) from C1, the percentage of larvae with increment widths similar to those of the survivors (Fig. 6B) increased shortly after hatching, reaching between 35% and 49% of the larvae, but then gradually decreased over time thereafter. Among the different OPT subgroups (100%, 75%, 50%, 25%), only the top 25% fastest-growing group reached increment 9, with more than 10% of its larvae displaying increment widths comparable to those of the surviving larvae. This suggests that even among the fastest-growing individuals in the initial cohort, fewer than 10% were in suitable physical condition to survive.

Fast-growing larvae, such as those in the 25% group, would have a higher likelihood of survival due to their higher growth rates (Fortier and Leggett, 1985; Aoki and Oozeki, 2007). Nevertheless, only a small proportion of the fastest-growing larvae within the optimum growth group were able to reach the MIW observed in the survivors (Fig. 6C), highlighting the narrow window of growth potential conducive to survival. Even among the best performers, only a small fraction is fit to withstand early life pressures. These findings underscore the disproportionate contribution of a minority of larvae to potential recruitment and the decisive role of growth rate in larval survival (Tanaka et al., 2022).

## 5. Conclusions

This study has examined the growth patterns and survival dynamics of Southern bluefin tuna (*Thunnus maccoyii*) larvae within their only known spawning habitat in the IO. The critical importance of larval physical condition at the onset of flexion appears to significantly influence survival during subsequent developmental stages. Larval growth rates observed in this study were higher than those reported more than three decades ago, possibly driven by an increase in sea surface temperature and, simultaneously, by differences variations in the composition and availability of prey in their pelagic environment. However, the observed thermal increase, although currently favorable for larval SBT development, is approaching the known biophysical limits for *Thunnus* species. Under future climate change scenarios, further warming could compromise both reproductive success and larval viability, especially in the IO. The importance of maternal inheritance in fast-growing SBT larvae lies in its enhancement of survival probability by increasing resistance and enabling early access to larger prey. Survival depends on a growth-selective process, in which only a very small percentage of fast-growing individuals since the first few days post hatch have a chance at survival. Overall, our results emphasize the need to continue and expand studies on the early life stages of SBT, especially in the context of continued warming of the IO and other future climate change impacts. Understanding how these conditions affect larval growth, survival, and selective processes during critical early life stages is fundamental to forecast recruitment variability and to inform sustainable management strategies for this ecologically and economically valuable species.

## Acknowledgements

We deeply acknowledge the dedication shown by the R/V Roger Revelle captain, crew and science party and the scientific contributions which went above and beyond during the RR2201 cruise. This study was supported by U. S. National Science Foundation awards OCE-1851558 (M.R.L.) and OCE-1851558 (D.D. and E.M.), and by PID2021-122862NB-100 INDITUN project from the Ministry of Science, Innovation and Universities (MICINN) of the Spanish government (R.L-C) and contributes to the Second International Indian Ocean Expedition (IIOE-2 endorsed project EP046). Seawater, zooplankton and larval fish samples were collected under Australian Government permit AU-COM2021-520 and Australian Marine Parks permit PA2021-00062-2 issued by the Director of National Parks, Australia. Views expressed in this publication do not necessarily represent those of the Director of National Parks or the Australian Government.

## Data Availability Statement

The data that support the findings of the present study are available from the first/corresponding author, R. Borrego-Santos, upon reasonable request

## Author Statement

R.B.-S., R.L.-C., J.M.Q., R.S., D.D and M.R.L conceived the study; R.B.-S. and F.A. analyzed environmental variables; R.B.-S., R.L.-C., J.M.Q. and E.M. collected and/or analyzed fish samples; R.B.-S., J.M.Q and E.M analyzed otolith microstructures; J.M.Q., R.B.-S., R.L.C., E.M., I.R., F.J.A. analyzed the data and/or discuss de results. All authors provided comments and edits on the manuscript.

## Declaration of Generative AI

Generative AI and AI-assisted technologies were solely employed during the writing process to improve the clarity and language quality of the manuscript.

